# Chromosome-scale *Salvia hispanica* L. (Chia) genome assembly reveals rampant *Salvia* interspecies introgression

**DOI:** 10.1101/2024.06.14.598901

**Authors:** Julia Brose, John P. Hamilton, Nicholas Schlecht, Dongyan Zhao, Paulina M. Mejía-Ponce, Arely Cruz Pérez, Brieanne Vaillancourt, Joshua C. Wood, Patrick P. Edger, Salvador Montes-Hernandez, Guillermo Orozco de Rosas, Björn Hamberger, Angélica Cibrian Jaramillo, C. Robin Buell

## Abstract

*Salvia hispanica* L. (Chia), a member of the Lamiaceae, is an economically important crop in Mesoamerica, with health benefits associated with its seed fatty acid composition. Chia varieties are distinguished based on seed color including mixed white and black (Chia pinta) and black (Chia negra). To facilitate research on Chia and expand on comparative analyses within the Lamiaceae, we generated a chromosome-scale assembly of a Chia pinta accession and performed comparative genome analyses with a previously published Chia negra genome assembly. The Chia pinta and negra genome sequences were highly similar as shown by a limited number of single nucleotide polymorphisms and extensive shared orthologous gene membership. There is an enrichment of terpene synthases in the Chia pinta genome relative to the Chia negra genome. We sequenced and analyzed the genomes of 20 Chia accessions with differing seed color and geographic origin revealing population structure within *S. hispanica* and interspecific introgressions of *Salvia* species. As the genus *Salvia* is polyphyletic, its evolutionary history remains unclear. Using large-scale synteny analysis within the Lamiaceae and orthologous group membership, we resolved the phylogeny of *Salvia* species. This study and its collective resources further our understanding of genomic diversity in this food crop and the extent of inter-species hybridizations in *Salvia*.

**PLAIN LANGUAGE SUMMARY:** Chia pinta is an economically important crop due to the high fatty acid present in the seeds. There are multiple types of Chia based on the seeds color including mixed which and black (Chia pinta), black (Chia negra), and white (Chia blanca). We generated a genome assembly of Chia pinta and compared it to existing genome assemblies. While the assemblies are highly similar there are key differences in terpene synthase composition between Chia pinta and Chia negra. We also sequenced 20 other Chia accessions with different seed color and geographic origin to determine a population structure within Chia. We generated genomic resources to further our understanding of this food crop.

**ABBREVIATIONS:** BGC Biosynthetic gene cluster

BUSCO Benchmarking Universal Single Copy Orthologs GO Gene ontology

SNP Single nucleotide polymorphism TIR Terminal inverted repeat

TPS Terpene synthase

WGS Whole genome shotgun

## INTRODUCTION

Chia (*Salvia hispanica* L.) belongs to the largest genus within the Lamiaceae containing approximately 980 species (Hu et al., 2018). Chia is a notable and economically important species within the *Salvia* genus attributable to the high nutritional value of its seeds which contain 16-26% protein, 23-41% fiber, and 20-34% polyunsaturated fatty acids, of which, 60% is *α*-lineolic acid (Muñoz et al., 2013). Historically, Chia was the third most economically important crop in Mesoamerica, only behind maize and amaranth, due to its use in religious practices and as a medicine (Valdivia-López & Tecante, 2015). The medicinal properties of Chia include treatments for gastrointestinal, respiratory, urinary, obstetrics, skin, central nervous, and ophthalmologic issues (Cahill, 2003). The traditional uses of Chia revolve around religious practices which contributed to the decrease of Chia prominence and cultivation in the 15^th^ century following the invasion by conquistadors (Cahill, 2003). Chia was introduced to Spain where it was named by Linnaeus as *Salvia hispanica* referencing the presumed origin of Spain (Baldivia, 2018). While Chia originated in present day Mexico and Guatemala, it has since been distributed throughout the world resulting in the emergence of diverse varieties (Cahill, 2004).

Chia varieties are characterized by their seed color and origin. The widely cultivated Chia blanca has a white seed coat while Chia negra has a black seed coat that can occur in wild and cultivated populations. Other seed coat colors include mixes of black and white seeds. Morphological characteristics distinguishing cultivated from wild accessions mirror traits observed in other domesticated species, such as decreased apical dominance, increased branching, increased seed size, decreased pubescence, increased florescence length determinism, increased anthocyanin pigmentation, variation in seed coat color and patterns, increased plant height, and closed calyxes (Cahill, 2004). While phenotypically distinct, dietary proteins are similar in wild and cultivated Chia accessions although wild accessions with higher levels of polyunsaturated fatty acids have been reported (Peláez et al., 2019).

Robust genomic resources for the Lamiaceae facilitate comparative genomic analysis. Within the Lamiaceae there are seven subfamilies with chromosome-scale genomes [Ajugoideae, Callicarpoideae, Nepetoideae, Lamiodeae, Scutellariodeae, and Tectonoideae] (Dong et al., 2018; Zhao et al., 2019a; b; Hamilton et al., 2020; He et al., 2022; Li et al., 2022; Shen et al., 2022; Sun et al., 2022; Pan et al., 2023). Current genomic resources for Chia include a genome assembly derived from an Australian black seeded variety (Chia negra; Wang et al., 2022), a white seeded variety (Chia blanca; Li et al., 2023), and a Mexican Chia (Alejo-Jacuinde et al., 2023) as well as transcriptomes constructed from wild and cultivated seeds (Peláez et al., 2019). Expanding the number and diversity of chia accessions with genome assemblies and sequence will facilitate our understanding of genetic diversity of this important crop as well as provide resources for more informed breeding programs. In addition to diversity within Chia, three other *Salvia* species occur in the same region in Mesoamerica (*Salvia uruapan* Fern., *Salvia tiliifolia* Vahl., and *Salvia polystachya* Ort.) that have similar uses as *S. hispanica* (Cahill, 2003). These species are challenging to distinguish from each other, but no reports indicate hybridization with *S. hispanica*. A phylogeny of *Salvia,* based on 91 nuclear genes, places Chia within *Salvia* sect. *Potiles* in a monophyletic clade (Lara-Cabrera et al., 2021). However, the *Salvia* genus has yet to be fully resolved and remains polyphyletic with *S. tiliifolia* being placed within two separate clades: the Angulatae and Polystachyae (Lara-Cabrera et al., 2021). Therefore, additional phylogenetic analyses are necessary to achieve a comprehensive resolution of the *Salvia* genus.

In this study, we report on the genome sequence of a Chia pinta accession, comparative analyses with published Chia genomes, and analysis of genetic diversity in a set of 20 Chia accessions revealing population structure between domesticated and wild Chia species and evidence of interspecies hybridization of *S. tiliifolia* with Chia.

## RESULTS AND DISCUSSION

### Chia Genome

We selected a Chia pinta accession from Acatic, Jalisco, Mexico that produces mixed color seeds and is grown as a superfood source throughout Mexico. Using 5.7 million PacBio long reads (36.5 Gb) representing ∼100x coverage of the predicted ∼355 Mbp Chia genome (Wang et al., 2022), we assembled the Chia pinta (2n=2x=12) genome using Canu (Koren et al., 2017). Whole genome shotgun (WGS) reads were used to generate a k-mer (k=21) distribution profile using GenomeScope indicating an estimated genome size of 338 Mbp with 62.6% unique kmers and 0.5% heterozygosity. The initial Canu assembly was error corrected using the raw PacBio reads using Arrow (Pacific Biosciences) followed by three rounds of error correction with the Illumina WGS reads using Pilon (Walker et al., 2014). The error-corrected assembly consisted of 2,094 contigs with a total length of 425.14 Mbp, which is substantially larger than the previously estimated genome size. Haplotigs were removed from the assembly using purgeHaplotigs (Roach et al., 2018) (-a = 50%) with an output consisting of “primary contigs” representing the putative haploid genome sequence, “haplotigs” containing diverged haplotypes, and “artefacts” representing contigs with very low or extremely high read coverage. Following removal of haplotigs, the “primary contigs” size decreased from 425 Mbp to 343 Mbp (Table 1). Manual examination of Chia vs. Chia self-alignments of contigs in the ‘purged assembly’ revealed five pairs of contigs that were putative residual haplotigs. Removal of these contigs resulted in a ‘purged assembly’ containing 407 contigs with an N50 contig length of 1.5 Mbp and a total size of 343.2 Mbp. The distribution of k-mers from WGS reads in the final assembly was examined using KAT (Mapleson et al., 2017) revealing a single peak indicating a haploid assembly with few retained haplotigs.

**Table 1.** Chia pinta Genome Assembly Metrics.

|  | **Input assembly** | **Purged Assembly** | **Final Chromosome-scale Assembly** |
| --- | --- | --- | --- |
| **Number of Contigs/**<br>**Chromosomes** | 2,094 | 407 | 6 |
| **Total length (bp)** | 425,143,449 | 343,219,856 | 341,980,016 |
| **Maximum Contig Length (bp)** | 9,374,111 | 9,374,111 | 67,233,260 |
| **Minimum Contig Length (bp)** | 1,684 | 2,780 | 57,181,130 |
| **N50 Contig Length (bp)** | 1,150,825 | 1,506,829 | 62,351,092 |
| **Average Contig Length (bp)** | 203,029 | 858,050 | 56,996,669 |

Using Hi-C sequence data, the contigs were assembled into six pseudomolecules, consistent with the known chromosome number of Chia and the Chia negra Australian Black (hereafter Chia negra) genome assembly (Wang et al., 2022). The final Chia pinta genome assembly was 342 Mb final with an N50 of 62Mb, of which, 99.64% of the assembly was anchored to one of the six pseudochromosomes (Table 1). Metrics for the final chromosome assembly were calculated using only the six chromosomes. The GC content of the final assembly was 36.6% consistent with the previously published Chia negra genome (Wang et al., 2022). Alignment of Illumina WGS reads to the final assembly revealed 98.4% of the reads aligned to the genome, of which, 99.5% were properly paired. Alignment of RNA-seq reads from a diverse set of tissue types (leaf, inflorescence, stem, and root) showed an overall alignment rate between 93.7% and 96.0%. To confirm the quality of the Chia pinta assembly, we used Benchmarking Universal Single Copy Orthologs (BUSCO) (Simão et al., 2015) to determine the representation of conserved orthologs in the final assembly. In total, 97.4% of the BUSCO orthologs were complete with 86.6% as single copy, 10.8% duplicated, 0.7% fragmented, and 1.9% missing. Overall, these results indicate a high-quality Chia pinta genome assembly.

### Repetitive Sequences and Transposable Element Annotation in the Chia pinta genome

Using *de novo* repetitive sequence identification with RepeatModeler coupled with sequences from the Viridiplantae RepBase, RepeatMasker masked 46.8% of the Chia pinta genome. With respect to transposable elements, retroelements were the dominant sequence with 40,151 retroelements occupying 15.15% of the Chia pinta genome while DNA transposons (36,807 elements) accounted for 4.86%. Unclassified interspersed repeats represented the largest number of elements with 378,795 or 26.11% of the genome. The remaining repetitive elements included rolling circle, small RNA, satellites, simple repeats, and low complexity sequences make up less than 1% of the genome.

The Extensive *de-novo* TE Annotator (EDTA) was used to annotate the Chia pinta genome for transposable elements revealing 314,306 elements spanning 149,780,410 bp (43.64%) of the Chia pinta genome. Long terminal repeats comprise 21.33% of the genome, of which, 5.7% were *Copia* elements and 11.45% were *Gypsy* elements; unknown long terminal repeats comprise 4.13% of the genome. Terminal inverted repeat (TIR) sequences represent 20.01% of the genome with the largest portion (12.09%) belonging to Tc1_Mariner family. The remaining TIRs are PIF_Harbinger (3.26%), hAT (2.32%), Mutator (1.80%), and CACTA (0.54%). Helitrons are non-terminal inverted repetitive elements and comprise 2.3% of the genome.

### Annotation of the Chia Pinta Genome

We annotated the Chia pinta genome for protein-coding genes resulting in 59,062 working gene models corresponding to 41,279 loci (Table 2). Working gene models had an average transcript length of 3.1 kbp, coding sequence (CDS) length of 1,217 bp, exon length of 279 bp and intron length of 240 bp. Working gene models exhibited an average of 5.8 exons, with 13.6% of transcripts being single-exon genes. The high confidence model set, a subset of the working set which have expression and/or protein evidence, contains 53,053 gene models representing 35,480 loci (Table 2). The high confidence set has an average transcript length of 3.3 kbp, exon length of 226 bp, intron length of 244 bp, and 6.1 exons per model; 6,105 gene models are single exon models. We selected the longest model as a representative for each gene locus from the working and high confidence model sets. With respect to BUSCO representation, the high confidence representative models are 94.8% complete, of which, 84.8% are complete and single copy while 10% are complete and duplicated; 1.9% are fragmented and 3.3% are missing. For the working representative models, 95.7% are complete with 85.5% complete and single copy and 10.2% complete and duplicated; 1.7% fragmented and 2.6% missing. Overall, the BUSCO results indicate a robust annotation of the Chia pinta genome.

**Table 2.** Chia pinta Genome Annotation Metrics.

|  | High<br>Confidence<br>Model Set | High<br>Confidence<br>Representative<br>Model Set | Working<br>Model<br>Set | Working Model<br>Representative<br>Set |
| --- | --- | --- | --- | --- |
| **Number of Gene Models** | 53,053 | 35,480 | 59,062 | 41,279 |
| **Number of Loci** | 35,480 | 35,480 | 41,279 | 41,279 |
| **Average Transcript Length (bp)** | 3,300.5 | 2,889.0 | 3,104.3 | 2,661.1 |
| **Average CDS Length (bp)** | 1,283.4 | 1,196.6 | 1,216.8 | 1,109.9 |
| **Average Exon Length (bp)** | 280.2 | 283.7 | 279.1 | 280.8 |
| **Average Intron Length (bp)** | 244.2 | 229.8 | 239.8 | 225.2 |
| **Average No. Exons per Model** | 6.1 | 5.3 | 5.8 | 4.9 |
| **Single Exon Transcripts** | 6,105 | 6,043 | 8,062 | 7,999 |

### Comparative Analyses of Chia Genome Assemblies

There are currently three published long-read, chromosome-scale Chia genome assemblies: Chia blanca (Li et al., 2023), Chia negra (Wang et al., 2022), and Mexican Chia (Alejo-Jacuinde et al., 2023). BUSCO analysis of all three published Chia genomes revealed that all of these assemblies were high quality and with robust gene annotation datasets. Syntenic orthologs (syntelogs) were identified between all four assemblies revealing a high degree of synteny between these genome assemblies (Figure 1) with limited disruptions that may be due to assembly artifacts in the various genome assemblies. Due to the high degree of similarity between the four Chia genomes, we performed detailed comparisons of our Chia pinta genome to the chromosome-scale black seeded Chia negra in which 73.62% of the genes were colinear within 1,178 syntenic blocks (Figure 1). Chia negra is a 344Mb genome assembly with 99.05% anchored on to chromosomes and 3.3Mb unanchored (Wang et al., 2022) with 428 gaps, amounting to a total of 191.2 kbp Ns. A total of 1,278,367 Single Nucleotide Polymorphisms (SNPs) were identified between the Chia negra and Chia pinta genomes that were distributed throughout the genome with 10.0% (127,210) residing in genic regions, 75.6% (967,385) in intergenic regions, and 14.4% (184,772) within intronic regions of the Chia pinta genome.

**Figure 1.**
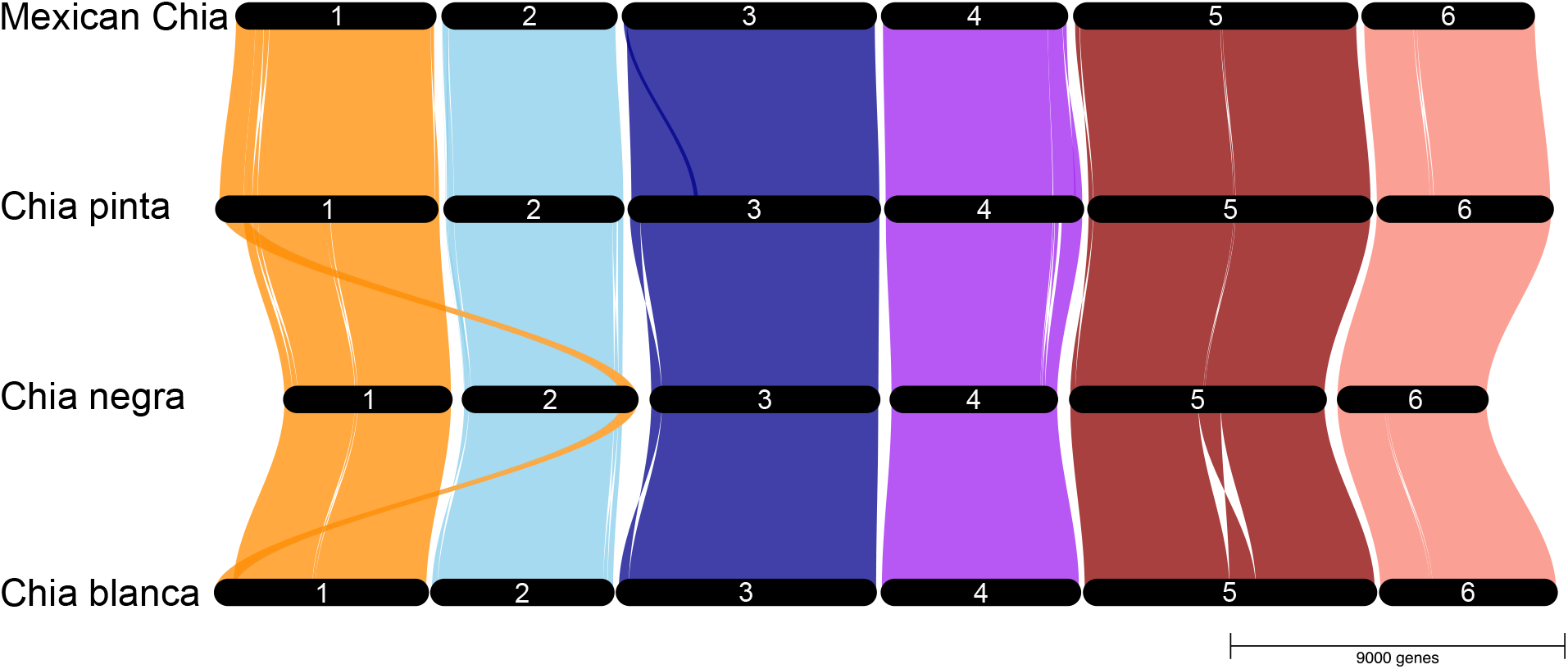
Synteny of the Chia genomes. The top track is the Mexican Chia genome (Alejo-Jacuinde et al., 2023), the second track is the Chia pinta genome reported in this study, the third genome is Chia negra (Wang et al., 2022), and the bottom track represents the Chia blanca genome (Li et al., 2023). The ribbons indicate syntenic blocks between the genomes identified using GENESPACE (v.1.1.10;Lovell et al., 2022).

Using Orthofinder with the predicted proteomes of both Chia pinta and Chia negra, we identified 20,580 orthogroups, of which, 358 orthogroups (2,738 genes) were unique to Chia pinta while 462 orthogroups (1,458 genes) were unique to Chia negra. Gene ontology (GO) enrichment of the genes unique to Chia pinta revealed differences in certain biological process, cellular components, and molecular function ontologies. Of particular interest was the enrichment of the GO terms “defense response”, and “diterpenoid biosynthetic process” with 45 terpene synthases identified in the GO terms “diterpenoid biosynthetic process” and “terpene synthase activity”.

BLASTP was used to search all representative proteins in Chia pinta and Chia negra against a collection of known terpene synthases (TPSs). TPSs greater than 350 amino acids were used to create a phylogeny to determine the relationships among the TPSs. After filtering, a total of 111 TPSs in Chia pinta and 53 in Chia negra were identified. To confirm that this is not due to annotation errors, Chia pinta TPS transcript sequences were used in a BLASTN search against the Chia negra genome; no additional terpene synthases were identified in Chia negra indicating these sequences are absent in the Chia negra genome assembly. A phylogeny was constructed with putative TPS protein sequences from Chia pinta, Chia negra, and functionally characterized TPSs to assign Chia TPSs to closest known functionally characterized TPSs. Despite GO enrichment annotation of ‘diterpenoid biosynthetic process’, most enriched TPSs are within the TPS-a and to a lesser degree TPS-b subfamilies which produce sesqui-and monoterpenes, indicating an expansion of volatile terpenes. The discrepancy on the GO terms claiming diterpenoid processes yet finding sesqui-and monoterpene synthases can be explained by GO enrichment often misannotated TPSs as diTPSs.

The TPS-a subfamily contains 56 putative TPSs in Chia pinta and only four in Chia negra. Of the 56 putative Chia pinta TPSs, 38 were found to enriched relative to Chia negra. The enriched TPSs reside in clades that do not contain a Chia negra TPS. To further understand the genomic context of the enriched TPSs, biosynthetic gene clusters (BGCs) membership and synteny were used. There are 16 BGCs containing TPSs in Chia pinta present on chromosomes 1, 2, 3, 4, and 6. Notably, six of these BGCs contain 23 out of the 56 Chia pinta specific TPS-a subfamily genes (Figure 2). This coincides with the expansion of the TPS-a subfamily in Chia pinta. All Chia pinta enriched TPS-a BGCs contain syntenic genes between Chia pinta, Chia negra, and *S. miltiorrhiza* (Figure 2). However, Chia pinta only shares one syntenic TPS with Chia negra and three syntenic TPSs with *S. miltiorrhiza.* Many of the TPSs present in Chia pinta’s BGCs appear to be tandem duplications, most notably in the teal and green BGCs (Figure 2). However, some of the TPSs present in the green BGC are less than 350 amino acids indicating they may be truncated.

**Figure 2.**
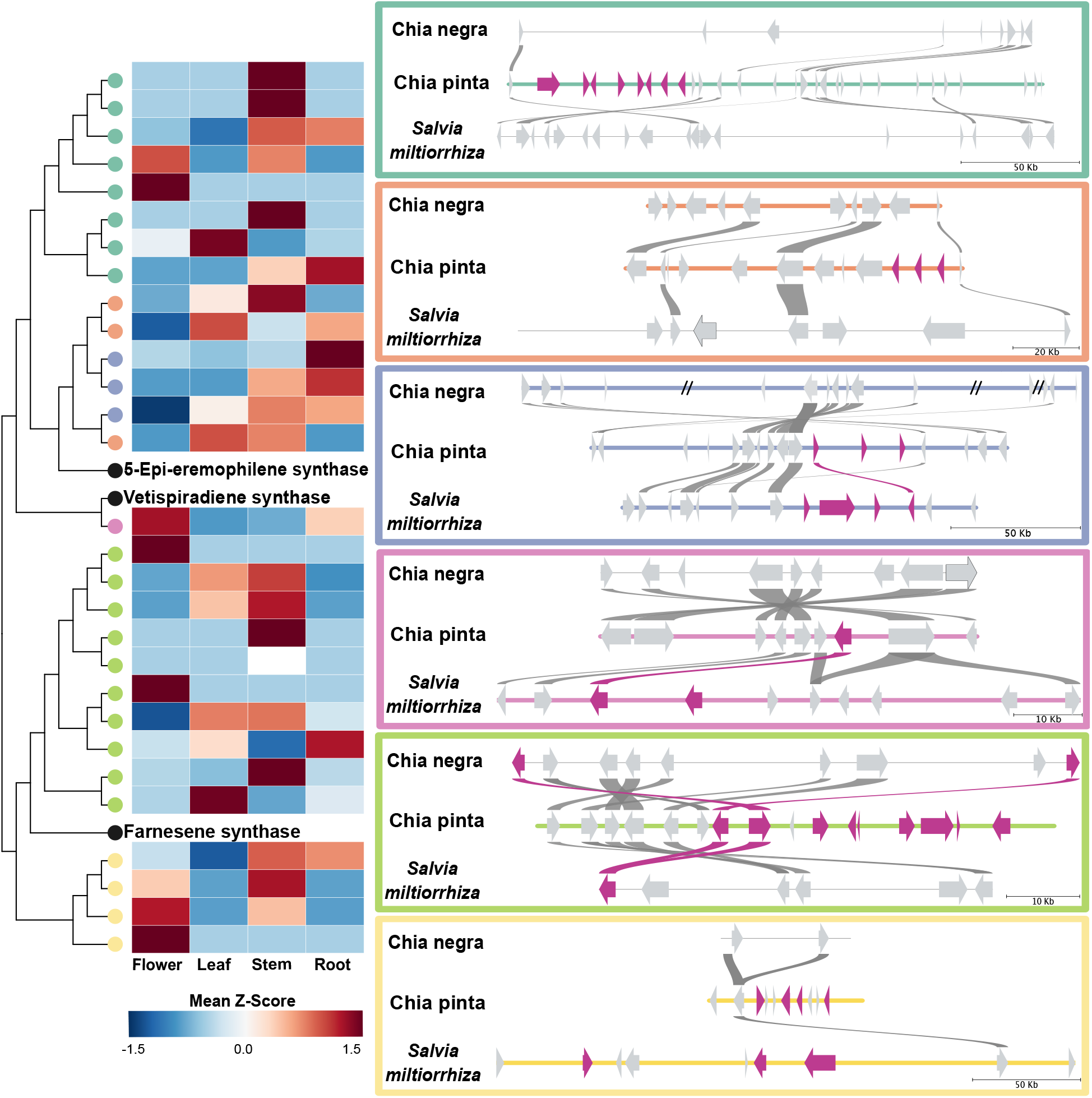
Chia pinta TPS-a Biosynthetic Gene Cluster Expression and Synteny. A phylogeny of the Chia pinta terpene synthase (TPS-a) genes present in biosynthetic gene clusters (BGCs) with representative functionally characterized reference TPSs is shown. The Chia pinta phylogeny was generated using RAxML (v8.2.12; Stamatakis, 2014). The heatmap of gene expression was constructed from flower, leaf, stem, and root tissue using expression values generated by Cufflinks (v.2.2.1; Roberts et al., 2011) with z-scores range from-1.5 to 1.5. Chia pinta genes (circles on the phylogeny) are colored by their respective BGC and correspond to the outlined syntenic BGCs; genes in black are known TPS. Biosynthetic gene clusters (BGCs) were identified by PlantiSmash (Kautsar et al., 2017) with boxes colored to match the clades in the phylogeny. Syntenic regions were determined using MCScanX (Wang et al., 2012) between Chia pinta, Chia negra, and *S. miltorrhiza*. Synteny is indicated as lines between the genes (arrows). The color of the gene and syntenic line is determined by the presumed identity assigned by PlantiSmash where hot pink indicate TPSs; slashes through the line indicate gaps in the assembly. Grey genome lines indicate that it is not a TPS BGC.

The origin and expansions of TPS-a genes were examined through synteny with *S. miltiorrhiza*. Two separate BGCs, purple and orange, contain paralogous TPSs yet are in distinct syntenic blocks (Figure 2). Work in *S. miltiorrhiza* characterized orthologs of these genes (89% identity) as (-)-5-epi-eremophilene synthases in which three TPSs (*SmSTPS1*, *SmSTPS2*, and *SmSTPS3*) had differential gene expression yet identical biochemical activity (Fang et al., 2017). The purple BGC contains one TPS that is a syntelog of *SmSTPS1*, but there are no syntelogs of *SmSTPS2* or *SmSTPS3* (Figure 2) suggesting that a single gene was maintained and was tandemly duplicated or that structural rearrangements occurred disrupting synteny with *SmSTPS2* or *SmSTPS3*. The orange BGC contains TPSs that are equally related to *SmSTPS1* but are not syntenic with the *S. miltiorrhiza* SmSTPS cluster. Instead, the homologs have moved into a different syntenic block entirely. Additionally, there is a notable difference in gene expression profiles of the purple and orange BGCs with the orange BGC largely expressed in the leaf and stem whereas the purple clade has its highest expression in roots amongst the different paralogs (Figure 2). This may exemplify how a BGC can evolve by duplication and subfunctionalization resulting in distinct spatial gene expression patterns. The teal and yellow BGCs indicate that there are no syntenic TPSs in *S. miltiorrhiza*. The minor enrichment in TPS-b genes present in Chia pinta is largely due to expansion of a single clade. The closest functionally characterized enzyme to this expanded clade was and (–)-exo-α-bergamotene synthase, having between 62-67% identity for this clade.

Finding such a large difference in TPS-a abundance and identifying many of them within BGCs between Chia pinta and Chia negra further supports the diversity that exists not just within the *Salvia* genus, but even within Chia accessions. One potential source of the TPS expansion could be due to sequencing gaps in the Chia negra genome assembly. Specially, there are gaps in the purple BGC region of the Chia negra genome sequence. Therefore, these TPSs could be present within the species, but were not captured by the genome assembly. However, for the remaining five BGCs there are no assembly gaps in the Chia negra genome assembly and when the predicted transcripts for the TPSs were searched against the Chia negra genome, there were no hits for these regions. To determine if the TPSs are unique to Chia pinta, we examined the BGCs for syntelogs in the two other long-read Chia genome assemblies. The teal, orange, pink, green, and yellow BGCs contain syntelogs in Chia pinta, Chia blanca, and Mexican Chia whereas the purple BGC contains only syntelogs between Chia pinta and Mexican Chia. Thus, diversity in TPSs is present between Chia accessions suggesting variation in terpenoid profiles that may be associated with local adaptation.

### Lamiaceae Phylogeny and Gene Family Expansions

To determine the evolutionary relationships of Lamiaceae species with Chia pinta, a species phylogeny was constructed using high-quality available genome sequences from 23 species from seven tribes in the Lamiaceae (Figure 3). Using the multiple sequence alignment option in Orthofinder, 923,746 genes were assigned to orthogroups. As shown in Figure 3, the Nepetoideae tribe is sister to Ajugoideae, Lamiodeae, and Scutellariodeae, the Callicarpoideae and Tectonoideae are sister to all other species, and the Premnoideae is sister to all other subfamilies. The relationships between the tribes in this genome-derived tree differs from a published phylogeny derived from 520 single copy transcripts (Godden et al., 2019) in which the Nepetoideae is sister to Ajugoideae, Lamiodeae, Scutellariodeae, Premnoideae, and Tectonoideae. The topology difference between these two phylogenetic estimates could be due to a combination of species sampling and data quality differences.

**Figure 3.**
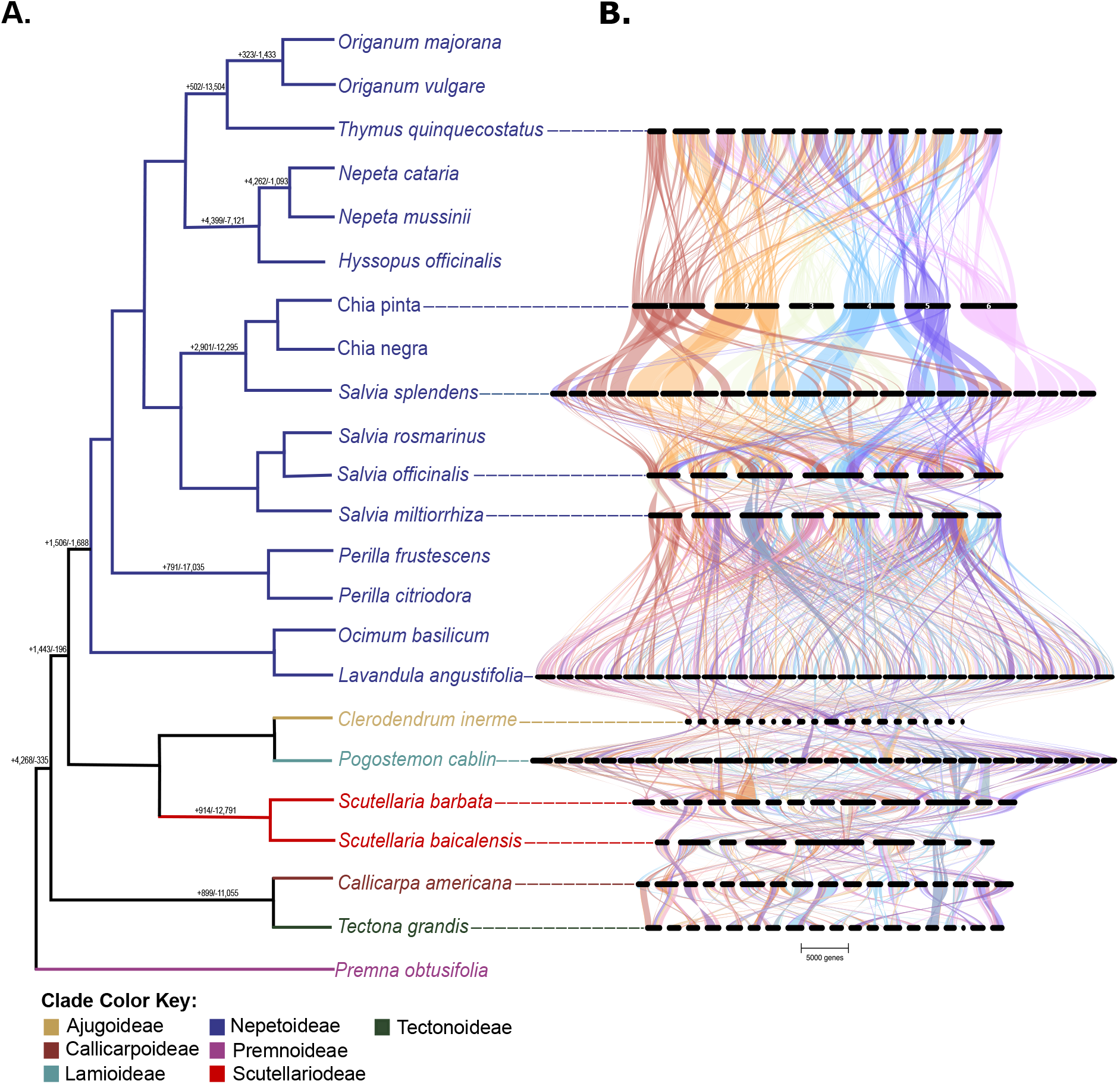
Lamiaceae phylogeny and synteny. A. A species phylogeny was generated using OrthoFinder (v.2.5.4;Emms & Kelly, 2019) using publicly available chromosome-scale Lamiaceae genomes. Numbers on branches indicated with (+) are gene family expansions and (-) are gene family contractions using CAFE (v.4.2.1; Han et al., 2013). **B.** The GENESPACE (v.1.1.10; Lovell et al., 2022) syntenic map of orthologous regions within chromosome-scale Lamiaceae genome assemblies are shown using the Chia pinta as the reference genome. Chromosomes are scaled by their physical length.

Gene expansions and contractions of single copy orthologs throughout the Lamiaceae were identified using CAFE (Figure 3A) and placed on the species tree phylogeny revealing large expansions and contractions throughout the Lamiaceae. The node branching of the Nepetoideae indicates a gene family expansion of 1,506 genes and contraction of 1,688 genes. The branch point from *S. hispanica* and *Salvia splendens* reveals 2,901 gene expansions and 12,295 gene contractions indicating substantial differences within the *Salvia* genus.

Synteny between genomes serve as a tool for examining evolution reflecting ancestral conservation of gene order. Using Chia pinta as the reference genome, we examined synteny within 11 chromosome-scale assemblies, spanning six tribes of the Lamiaceae family, revealing extensive conservation among the genomes (Figure 3B). In total, 182 Chia pinta genes were found to have a one-to-one syntenic relationship across all 11 species.

The polyphyletic nature of *Salvia* is highlighted by orthogroup membership. Of the 39,379 orthogroups containing 211,888 genes there were 12,987 orthogroups, containing 165,520 genes, in common among all *Salvia* (Figure 4A). The next highest number of orthogroups are unique to *S. rosmarinus* closely followed by *S. officinalis* and then *S. splendens* (Figure 4A). We also performed syntenic analyses between the genomes of four *Salvia* species to further our understanding of the species relationship in this polyphyletic genus. As expected, Chia pinta shares extensive synteny with other *Salvia* species (Figure 4b). *S. splendens* is reported to be a tetraploid (Jia et al., 2021). Based on orthogroup membership, 25% (4,684) of orthogroups shared by *S. splendens* and Chia pinta contain two *S. splendens* genes for each Chia pinta gene. This pattern reflects that *S. splendens* is a tetraploid and Chia pinta is a diploid. There are also two syntenic blocks in *S. splendens* for each block within Chia pinta, the syntenic blocks exist across four chromosomes in *S. splendens* (Figure 4b and 4c). It has been reported that there is a single shared whole genome duplication between Chia pinta and *S. splendens* and an additional duplication just in *S. splendens* (Jia et al., 2021; Wang et al., 2022). Therefore, the four unique chromosomes syntenic to a single chromosome in Chia pinta could be due to chromosomal fusions in Chia pinta or chromosomal fissions in *S. splendens.* Within the *Salvia* genus there are large regions of fragmented synteny between Chia pinta and *S. officinalis* as well as between *S. splendens* and *S. officinalis.* The fragmentation could be present due to different ancestry of Chia pinta and *S. officinalis*. As *Salvia* is a polyphyletic genus (Lara-Cabrera et al., 2021), this could be indicative of how distantly related these two species are. An alternative hypothesis is that they share a common ancestor, but the divergence time between species is so long that conserved genetic regions have been differentially fractionated (i.e. unique gene loss patterns). This is consistent with the large gene family expansions and contractions in the node that splits the *Salvia* species.

**Figure 4.**
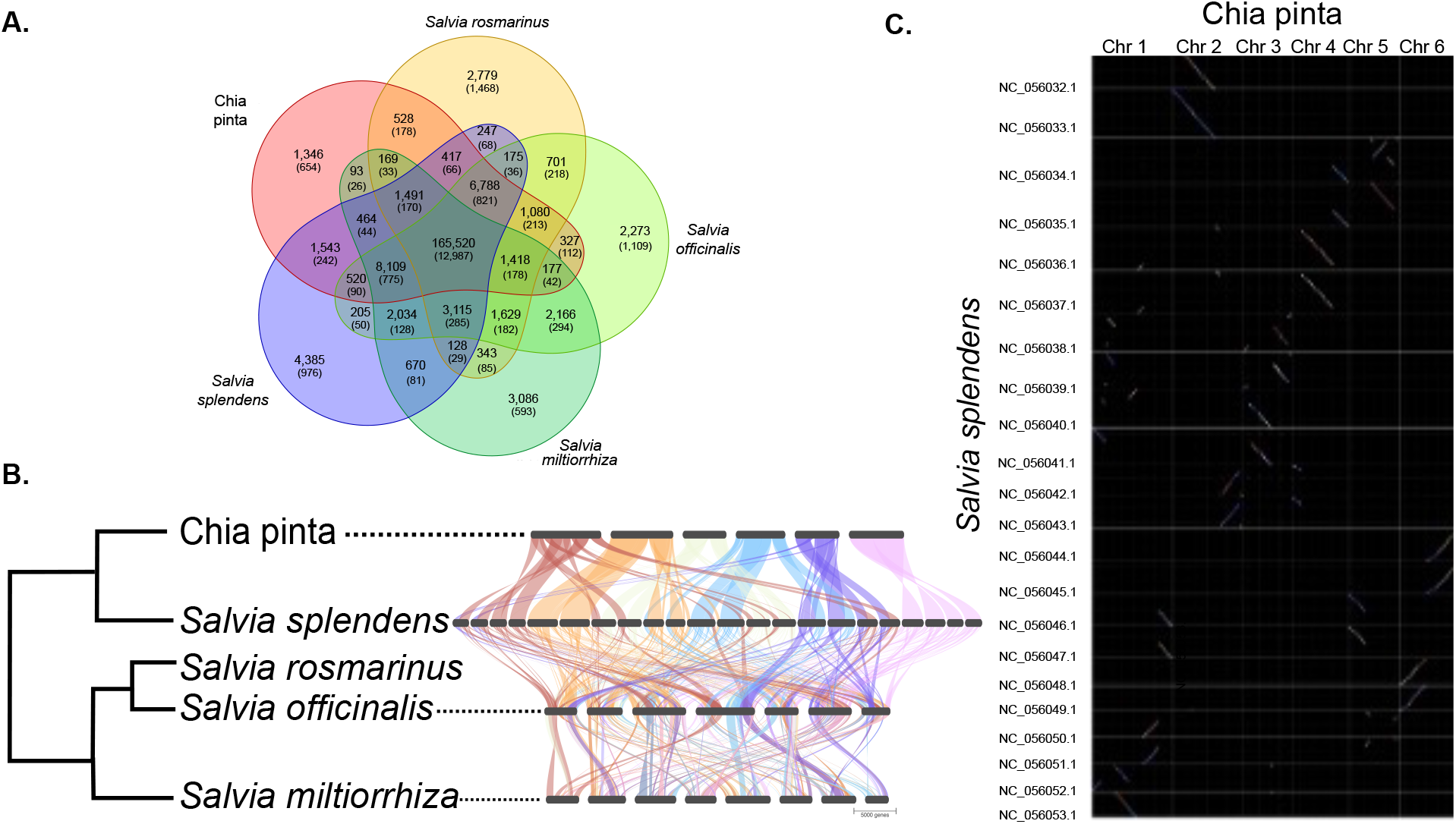
*Salvia* gene orthology and synteny. A. *Salvia* orthogroup intersections between Chia pinta, *Salvia rosmarinus*, *Salvia officinalis*, *Salvia splendens,* and *Salvia miltiorrhiza* as determined by OrthoFinder (v.2.5.4; Emms & Kelly, 2019). Numbers of orthologous groups and genes in parentheses are reported. **B.** GENESPACE (v.1.1.10; Lovell et al., 2022) syntenic map of orthologous regions within chromosome-scale *Salvia* genome assemblies using Chia pinta as the reference genome. **C.** Synteny dotplot for the anchor genes between Chia pinta and *Salvia splendens* generated in GENESPACE (v.1.1.10; Lovell et al., 2022). Chia pinta includes 21,720 genes with BLAST hits. *Salvia splendens* includes 25,958 genes with blast hits.

### Population Structure of Chia

Seed coat color is a frequent descriptor for Chia accessions with Chia white seeded blanca varieties while Chia negra, Chia cualac, and Chia xonostli are predominately black-seeded (Figure 5a). Chia pinta seeds are a mix of both black and white seeds (Figure 5a). A diversity panel of 19 Chia accessions including wild and cultivated accessions along with two *S. tiliifolia* accessions with origins throughout Mexico was constructed and sequenced to reveal genetic diversity among accessions and provide insight into population structure of cultivated and wild Chia varieties. The percentage of reads aligned to the Chia pinta genome ranged from 95.5% to 97.7% for the *S. tiliifolia* samples and 96.3%-98.5% for the Chia varieties suggesting that the two species share substantial sequence similarity. Population structures were inferred with admixture with k=2 to k =13. Population structure admixture results suggested through the cross-validation error plot that there are two possible number of populations: four and nine as the local minima being at four and the global minima at nine in the cross-validation error plot.

**Figure 5.**
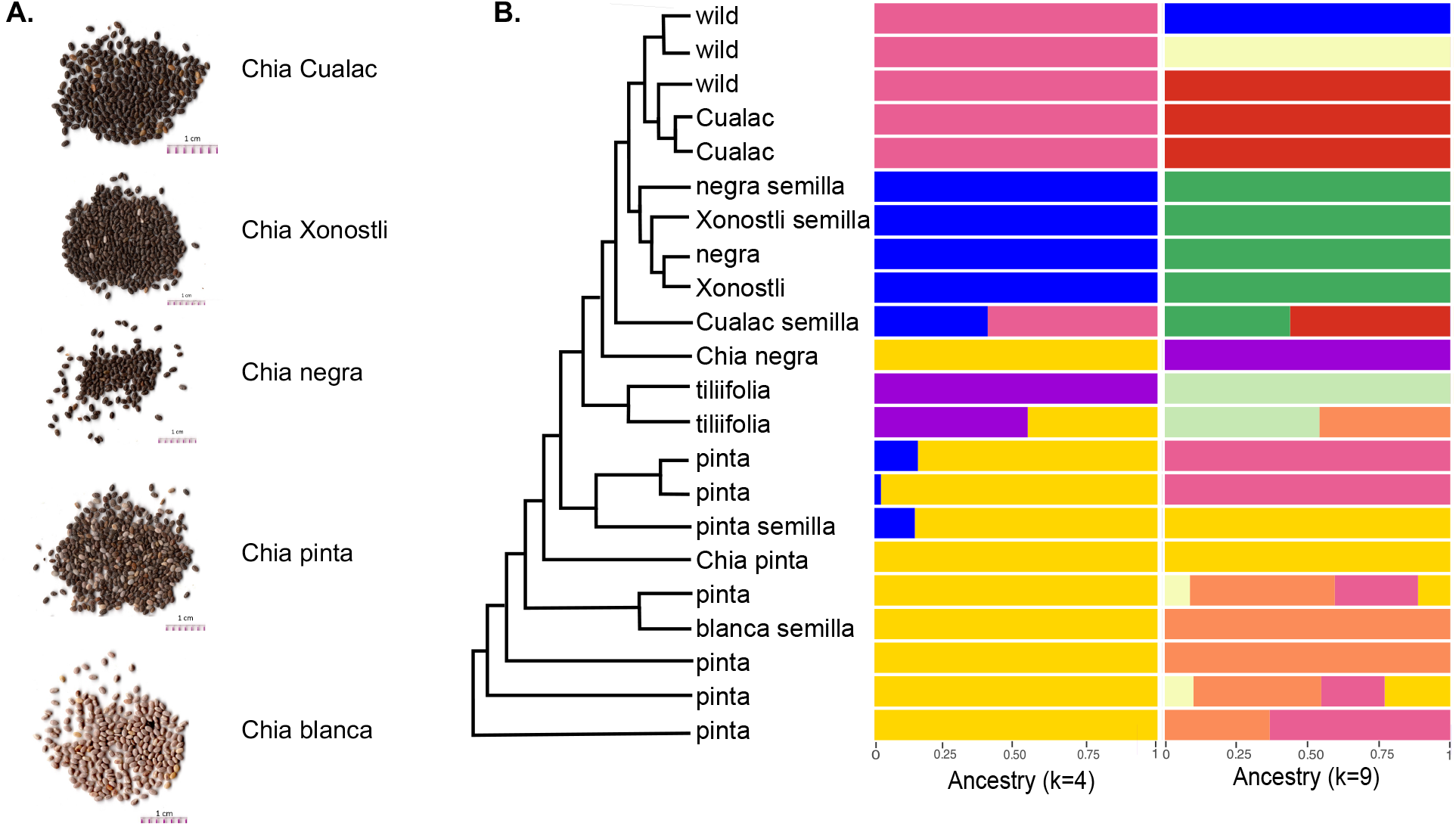
**Population structure of Chia**. **A.** Representative seed images of Chia varieties. **B.** SNP phylogeny was built using SNPhylo (v.20160204; Lee et al., 2014). Admixture (v.1.3.0; Alexander et al., 2009) population structure of 20 Chia accessions and 2 *Salvia tiliifolia* accessions was generated from 156,829 SNPs. Populations from the minima on the cross-validation plot was determined using k=4 and k=9.

Using a k=4, broad population groups are present that can be assigned to known categories of Chia: Chia pinta (yellow), *S. tiliifolia* (purple), Chia negra and Chia Xonostli (blue), and Chia Cualac (pink). The population structure indicates that the phenotypic and origin grouping reflects the genetic structure of the population. Chia pinta accessions are domesticated Chia varieties whereas Chia negra and Chia Xonostli are classified as wild due to their more open calyx and other wild traits. Chia negra is in the same population group with the less widely known Chia Xonostli which is similar to Chia negra yet categorized differently due to its domesticated traits. Historically, Chia Xonostli was found in the states of Jalisco, Guanajuato, Veracruz, and Hidalgo. Chia Cualac was reported to be semi-domesticated and forms their own group with some admixture from Chia Xonostli (Peláez et al., 2019). This follows the hypothesis that wild introgressions are present throughout the populations. One *S. tiliifolia* accession is admixed with Chia pinta. *S. tiliifolia* is nearly indistinguishable from Chia and is known to grow in the same areas as Chia pinta; thus, it is possible that these species hybridize and form a population of *S. tiliifolia* that is highly admixed with Chia pinta. Feral hybrid accessions could continue to evolve through hybridization with domesticated Chia yielding the admixture present within one accession of *S. tiliifolia* (Figure 5).

## CONCLUSIONS

In this study, a high-quality chromosome-scale genome assembly of Chia pinta was generated that allowed for additional genomic comparisons within the economically important crop including three other long-read, chromosome-scale Chia assemblies that showed extensive synteny among the genome sequences. Comparative genomic tools were used to determine differences within Chia accessions and throughout the Lamiaceae. Interestingly, Chia pinta was enriched in TPSs and contains novel TPSs compared to the Chia negra with some TPSs located within BGCs and syntenic with *S. miltiorrhiza*. Further examination of TPSs within BGCs among the four Chia genome assemblies revealed further diversification suggestive of variation in terpenoid biosynthesis among varieties. Through sequencing of a diversity panel, the population structure of Chia revealed introgression with other *Salvia* species.

## MATERIALS AND METHODS

### Plant materials

Different Chia varieties were collected throughout Mexico. Plants were grown in an experimental field in Celaya, Guanajuato, Mexico (20.578°, −100.822°) at the Instituto Nacional de Investigaciones Forestales, Agrícolas y Pecuarias (INIFAP).

### Nucleic acid isolation, library construction, and sequencing

For construction of a reference genome, DNA was isolated from medium-sized leaves from a mature plant (13.5 weeks old) of accession SM_ACJ2017 using a modified protocol from Doyle and Doyle (1987) and Healey et al. (2014). Large insert (>15kb, >20kb) PacBio libraries were made with the SMRTbell™ Template Prep Kit and sequenced on the PacBio Sequel platform at the University of Georgia, Georgia Genomics and Bioinformatics Core (GGBC, UG Athens, GA, RRID:SCR_010994). A whole genome shotgun library for reference error correction was prepared using the Illumina TruSeq Nano DNA Library Preparation Kit and sequenced in paired-end mode, 150 nt in length on a HiSeq 4000 at the Michigan State University Research Technology Support Facility (RTSF). Whole genome shotgun libraries for use in error correction and diversity panel variant analyses were constructed as described previously in Hardigan *et al*. 2016 (Hardigan et al., 2016) and sequenced at the Michigan State University RTSF in paired-end mode on a HiSeq4000 generating 150 nt reads. RNA was isolated from three biological replicates from a core set of tissues (leaf, inflorescence, lateral stem, secondary root) from the reference accession SM_ACJ2017 as described previously in Peláez *et al*. 2019 (Peláez et al., 2019). RNA-seq libraries were prepared using the Illumina TruSeq Stranded mRNA Library Preparation Kit and sequenced on an Illumina HiSeq 4000 generating 150 nt paired end reads for one replicate and 50 nt single end reads for the other two replicates; library preparation and sequencing were performed at the Michigan State University Research Technology Support Facility (RTSF). A Phase Genomics Proximo Hi-C library was prepared from Chia pinta leaf tissue and sequenced by Phase Genomics (Seattle, WA) on the NextSeq 500 generating paired end 150 nt reads.

### Chia pinta genome assembly

PacBio reads greater than 10 kbp (1.2 million reads, 21.6 Gb) were used to generate the initial assembly using Canu (v1.7; Koren et al., 2017) with a corrected ErrorRate of 0.15%. The initial assembly was polished with the raw PacBio reads using Arrow in the SMRT Analysis package (v5.0.1.9585; Pacfici Biosciences), followed by three rounds of error correction with 56 million Illumina WGS reads (150 nt paired-end WGS reads, 45X coverage) using Pilon (v1.22; Walker et al., 2014). Potential haplotigs were purged using purgeHaplotigs (v1.0.4; Roach et al., 2018) with the “maximum match score (-m)” of 500% and “-a = 50% “. Contigs were scaffolded to a chromosome scale assembly using Hi-C reads and Proximo pipeline with an input chromosome number of six by Phase Genomics (Bickhart et al., 2017). Scaffolded contigs were visualized with Juicebox (v1.9.8; Durand et al., 2016).

### Genome annotation

A custom repeat library (CRL) was generated using RepeatModeler (v2.0.1;Flynn et al., 2020) and protein coding genes were removed from the CRL using ProteinExcluder (v1.2; Campbell et al., 2014). The Viridiplantae RepBase repeats (v20150807) were then added to create the final CRL. The genome assembly was hard and soft masked using RepeatMasker (v4.1.0; Smit et al.) with the CRL with the parameters:-s-nolow-no_is. RNA-seq libraries were cleaned using Cutadapt (v2.9; Martin, 2011) (--times 2--minimum-length 100--quality-cutoff and then aligned to the genome assembly with HISAT2 (v2.2.0;Kim et al., 2019) (--max-intronlen 5000--rna-strandness RF –dta –no-unal). The RNA-seq alignments were then assembled into transcript assemblies using Stringtie (v2.1.1; Kovaka et al., 2019).

*Ab initio* gene models were predicted on the soft-masked genome assembly using the BRAKER2 pipeline (v2.1.5; Brůna et al., 2021) using the leaf RNA-seq library CHI_AA as a source for hints. The *ab initio* gene models were then refined using PASA2 (v2.4.1; Campbell et al., 2006) with the RNA-seq transcript assemblies as a source of transcript evidence to produce the working gene model set. High confidence gene models were selected from the working gene model set by first calculating working gene model abundances of the RNA-seq libraries for the working gene models with Kallisto (v0.46.0; Bray et al., 2016), then searching the working gene models against PFAM (v32.0; Mistry et al., 2021) with HMMER (v3.2.1; Mistry et al., 2013). Working gene models with a TPM >1 in at least one RNA-seq library or a non-transposable element related PFAM domain match and no partial or containing an internal stop codon were identified as high confidence gene models. Functional annotation was assigned to the working gene model by searching the protein sequences against the Arabidopsis proteome (TAIR10), PFAM (v32.0; Mistry et al., 2021) and the Swiss-Prot plant proteins (release 2015_08). Search results were processed in the same order and the function of the first hit encountered was assigned to the gene model. Repetitive elements were identified using EDTA (v2.1.0; Ou et al., 2019) with the parameters species set to “others” and step set to “all”.

### Genome quality assessment

Quality assessment of the genome assembly was performed by aligning WGS reads cleaned for low quality bases and adaptors using Cutadapt (v3.4; Martin, 2011) to the final assembly using BWA-mem (v0.7.16a; Li, 2013). Assemblathon.pl (https://github.com/KorfLab/Assemblathon/blob/master/assemblathon_stats.pl) was used to generate genome metrics. BUSCO (v3.1.0.Py3; Simão et al., 2015) embryophyta_odb10 was used to determine genic representation in the final assembly. Jellyfish (v.2.3.0; Marçais & Kingsford, 2011) with the option-m 21 was used to count kmers that were then visualized in GenomeScope (v2.0; Ranallo-Benavidez et al., 2020) with kmer length 21 was used to verify genome size and heterozygosity from the WGS reads from Chia pinta (CHI_AN). The Kmer Analysis Toolkit (v2.4.1; Mapleson et al., 2017) was used to examine the assembly for retained haplotigs. Synteny between the chia genome assemblies (Wang et al., 2022; Alejo-Jacuinde et al., 2023; Li et al., 2023) was analyzed using GENESPACE (v.1.1.10;Lovell et al., 2022). Syntenic comparison between Chia pinta and Chia negra was also performed using MCScanX (Wang et al., 2012).

### Lamiaceae phylogeny and comparative analysis

Publicly available genomes of *Callicarpa armericana* (Hamilton et al., 2020), *Cleorodendrum inerme* (He et al., 2022), *Hyssopus officinalis* (Lichman et al., 2020), *Nepeta cataria* (Lichman et al., 2020), *Nepeta mussinii* (Lichman et al., 2020), *Ocimum basilicum* (Bornowski et al., 2020), *Origanum majorana* (Bornowski et al., 2020), *Origanum vulgare* (Bornowski et al., 2020), *Perilla frustescens*(Zhang et al., 2021; Tamura et al., 2022), *Pogostemon cablin* (Shen et al., 2022), *Salvia miltiorrhiza* (Pan et al., 2023), *Salvia officinalis* (Li et al., 2022), *Salvia rosmarinus* (Bornowski et al., 2020), *Salvia splendens* (Jia et al., 2021), *Scutellaria baicalensis* (Zhao et al., 2019b), *Scutellaria barbata* (Xu et al., 2020), *Tectona grandis* (Zhao et al., 2019a), *Thymus quinquecostatus* (Sun et al., 2022)*, Lavandula angustifolia* (Hamilton et al., 2023) and *Premna obstusifolia* (He et al., 2022) were obtained and quality assessed using BUSCO (v5.5.0; Simão et al., 2015) embryophyta_odb10. Species with genome assembly complete BUSCO scores greater than 90% and annotation complete BUSCO scores greater than 80% were used in further comparative analysis. Orthogonal genes and species tree phylogeny were built using OrthoFinder (v.2.5.4; Emms & Kelly, 2019) with options-M msa-T raxml. The species tree output was covered into an ultrametric tree using the make_ultrametric command in OrthoFinder (v.2.5.4; Emms & Kelly, 2019). Branch lengths were rescaled using the *Premna obstusifolia* divergence date of 16.06 MYA retrieved from the TimeTree of Life resource (Kumar et al., 2022). Gene family expansions and contractions were identified using CAFE (v.4.2.1; Han et al., 2013) with the following scripts with default parameters: cafetutorial_report_analysis.py and cafetutorial_draw_tree.py. Syntelogs through the Lamiaceae were obtained for the chromosome scale assemblies within the Lamiaceae and visualized using GENESPACE (v.1.1.10; Lovell et al., 2022).

### Gene ontology term enrichment

Gene ontology (GO) terms were assigned to high confidence Chia pinta genes using InterProScan (v5.63-95.0; Jones et al., 2014). GO descriptions were added using the ontologyIndex package (Greene et al., 2017) and enrichment was calculated using the topGO R package (Alexa & Rahnenfuhrer, 2010). GO terms with an FDR adjusted p-value < 0.05 were considered significant.

### Terpene synthase identification

BGCs were identified in Chia pinta, Chia negra, and *S. miltiorrhiza* with PlantiSMASH (Kautsar et al., 2017). Enriched TPSs identified in the various BGCs were searched with NCBI BLAST the nonredundant protein database to identify the closest functionally characterized TPSs. To extract all TPSs from the genome, the high confidence representative protein models were blasted against a reference set of known TPSs enzymes representing TPSs across all subfamilies. The BLAST hits with an E-value 1E-5 or better were selected. These gene models were filtered to remove any sequences smaller than 350 amino acids to ensure a quality phylogeny and minimize pseudogenes. The final set of putative and reference TPS sequences were aligned using clustal omega (v1.2.4; Sievers et al., 2011). A phylogenetic tree of the alignment was built via RAXML (v8.2.12; Stamatakis, 2014) with the PROTGAMMA AUTO model, algorithm a, and 1000 bootstraps. Gene expression of terpene synthases was calculated using the single end RNA-seq libraries and Cufflinks (v.2.2.1; Roberts et al., 2011) with the options-b and-u to generate FPKM values for all Chia pinta genes. Orthologous genes from Chia pinta, Chia negra (Wang et al., 2022), Chia blanca (Li et al., 2023)and the Mexican Chia variety (Alejo-Jacuinde et al., 2023) were identified using OrthoFinder (v.2.5.4; Emms & Kelly, 2019) with options-M msa-T raxml.

### Population structure analysis

Whole genome shotgun reads from the diversity panel were cleaned using Cutadapt (v3.4; Martin, 2011) and aligned to the Chia genome using BWA-mem (v0.7.16a; Li, 2013). PicardTools (v2.20.8; Picard toolkit, 2019) commands SortSam, MarkDuplicates, BuildBamIndex, and CollectAlignmentSummaryMetrics were used to sort, convert files, and generate alignment metrics. The GATK (v4.1.2.0; Van der Auwera & O’Connor, 2020) HaplotypeCaller with default parameters was used to call variants. GenomicsDBImport with default parameters was used to merge the varieties into a single VCF file and genotyped using GenotypeGVCFs.Separated. SNPs were selected using the SelectVariants command. Hard filtering of the SNPs was performed using the parameters QD < 2.0, QUAL < 30.0, SOR > 3.0, FS > 60.0, MQ < 40.0, MQRankSum <-12.5, MQRankSum-12.5, ReadPosRankSum <-8.0. Additional filtering was performed using VCFTools (v0.1.16; Danecek et al., 2011) with filtering –freq2 and –max-alleles 2 to retain only bialleleic sites, minor allele frequency of 0.071,--max-missing 0.9,--minQ 30,--min-meanDP 15,--max-meanDP 39.

SNPs were called relative to the Chia negra reference genome (Wang et al., 2022) using nucmer from MUMmer (v4.0; Marçais et al., 2018) with the options –maxgap=2500,--minmatch=11, and--mincluster=25. SNPs were quality filtered using the delta-filter command in MUMmer with the-r flag. (v4.0; Marçais et al., 2018). The SNP set from the diversity panel and from the alignment of the two genome assemblies were combined and converted into bed format using PLINK 2.0 (v.alpha2.3; Purcell & Chang; Chang et al., 2015) resulting in 156,829 total SNPs. Population structure was inferred with Admixture (v.1.3.0; Alexander et al., 2009) and a SNP phylogenetic tree built with SNPhylo (v.20160204; Lee et al., 2014) using default parameters.

## ACKNOWLEDGMENTS

Funds for this study were provided by a grant to C.R.B. from the National Science Foundation Plant Genome Research Program (IOS-1444499), the Georgia Research Alliance, Georgia Seed Development, and the University of Georgia. JB was supported by Michigan State University.

## CONTRIBUTIONS

ACJ and CRB conceived of the study. SM-HGOR, PMM-P and ACP collected samples. PMM-P and ACP prepared materials. JB, JPH, NS, DZ, JCW and BV performed data analyses and drafted the manuscript. PPE, BH, ACJ, and CRB supervised and performed project administration. All authors approved of the manuscript.

## DATA AVAILABILITY STATEMENT

The raw sequence reads are available in the National Center for Biotechnology Information Sequence Read Archive under BioProject PRJNA744892. The genome assembly, annotation, and large data sets (genome assembly, genome annotation) reported in this study are available in Figshare via 10.6084/m9.figshare.24546049.

